# Reactivating aversive memories in humans: An EEG and Microstates study of post-retrieval processes

**DOI:** 10.1101/2025.11.03.686328

**Authors:** Luciano Cavallino, Luz Bavassi, Maria Eugenia Pedreira

**Author notes:** Corresponding author Maria Eugenia Pedreira, Ph.D. Laboratorio de Neurociencias de la Memoria, Instituto de Fisiología, Biología Molecular y Neurociencias (IFIBYNE, CONICET), Facultad de Ciencias Exactas y Naturales, Universidad de Buenos Aires. Av. Costanera Rafael Obligado, Ciudad Universitaria, Ciudad Autónoma de Buenos Aires, (C1428EGA), Buenos Aires, Argentina.

## Abstract

Although fear conditioning is one of the most widely used models to study anxiety in humans, much remains to be explored regarding its underlying neural correlates. In this study, we employed a three-day threat-conditioning protocol in which an angry face (conditioned stimulus, CS+) was paired with an aversive sound (unconditioned stimulus, US). Our main objective was to identify neural markers of post-retrieval processes triggered by the presentation of the CS+ alone, 24 hours after acquisition.

We recorded resting-state electroencephalographic (EEG) activity before and after memory reactivation in two groups of participants (females and males counterbalanced): a Reactivation group previously exposed to threat conditioning on Day 1 (n=26) and a Control group (n=26) with no prior conditioning, focusing on memory-related patterns in the frequency domain and microstate dynamics.

Our results confirmed successful conditioning and memory retention 48 hours later, as evidenced by implicit, declarative, and emotional responses. Regarding neural activity, we found lower beta-band activity (25–30 Hz) in central brain regions after reactivation, which may reflect enhanced post-reactivation processing of the threat conditioning memory, in contrast to participants who viewed the stimulus for the first time. We observed changes in microstates and a potential link between microstate C and conditioned threat memory.

These findings provide initial evidence for neural correlates of post-retrieval processes of an implicit threat memory offering potential relevance for understanding mechanisms implicated in anxiety and related disorders.

## Introduction

Understanding environmental threats serves as a critical survival mechanism, allowing organisms to associate previously neutral stimuli with aversive events. This leads to the formation of implicit memories that initiate defensive behaviors, thereby helping to prevent direct harm and enhancing survival rates^1^. However, when an organism fails to suppress threat responses in safe environments, these survival behaviors can become maladaptive, resulting in dysfunctional reactions. Many researchers propose that these mechanisms may contribute to anxiety disorders^2^. In this context, threat-conditioning paradigms offer a powerful experimental approach for investigating the neurobiological foundations of anxiety disorders. Foundational studies in animal models have identified the amygdala as a central hub for the associative learning of neutral cues and aversive outcomes^3–6^. In humans, threat conditioning demonstrates that skin conductance responses are indicative of autonomic nervous system activity^7^, acting as a reliable marker for the implicit expression of threat memories, even in the absence of conscious awareness of the underlying contingencies^8–10^. Clinical studies further illustrate that altered threat conditioning and extinction processes are characteristic of anxiety disorders, thus linking fundamental mechanisms of associative learning to psychopathology^11–13^. Collectively, this body of research has established threat conditioning as a cross-species model that connects conserved neural and physiological processes with maladaptive fear and anxiety.

The resurgence of experiments suggesting that specific cues can trigger the reactivation of memories has called into question the longstanding assumption that consolidated memories are permanently resistant to modification^14,15^. This has led to the development of the reconsolidation framework, which posits that the presentation of a reminder triggers reactivation and acts as a mechanism facilitating memory malleability. A closer examination of these protocols reveals that the effectiveness of these reminders relies on the presentation of partial information from the original task. In general, incomplete reminders are characterized by the omission of the expected outcome^16^. However, a deeper analysis of the reminder structures indicates that their effectiveness relies on providing cues that contain critical information, thereby prompting attempts at retrieval^16,17^. Taking this into account, the reactivation of a threat-conditioning memory typically involves reminders in which the unconditioned stimulus (US) is omitted. ^14,18^

Anxiety disorders involve physiological, emotional, and cognitive alterations. However, experimental paradigms typically focus on associative responses assessed through autonomic indices (e.g., heart rate variability, skin conductance) and declarative measures (e.g., contingency awareness). This emphasis neglects key components of threat processing, such as cognitive bias, defined as systematic deviations from rational judgment^19,20^. To bridge this gap, we developed a protocol that integrates physiological and declarative measures with tasks revealing cognitive bias, providing a novel tool to investigate cognitive disturbances associated with anxiety disorders^21^. More recently, we demonstrated that presenting a prediction error reminder (CS+ without US) 24 hours after conditioning led to the labilization of the previously formed memory in a threat conditioning paradigm^22^. This conclusion was based on findings that performing a highly demanding task immediately after memory reactivation eliminated the increased SCR to the CS+ during testing and abolished the cognitive bias effect. These results support the idea that the demanding task weakened the memory trace, but only when the memory had first been reactivated and rendered labile by the reminder.

Given that memory processes rely on neural communication and plasticity across a broad range of timescales, it has been proposed that oscillatory brain activity, ranging from milliseconds to seconds, carries information about higher-order cognitive processes, both in terms of amplitude dynamics and phase coherence^23^. For example, it has been demonstrated that decreases in amplitude within the 8–40 Hz frequency range play an active role in the formation and retrieval of long-term episodic memories^24^.

In fMRI studies, evidence has been found of increased spontaneous activity in specific brain regions, as well as greater functional connectivity when a fear reminder (CS + US) was presented, compared with a group that did not receive reinforcement, during post-stimulus resting-state evaluations^25^. Other studies showed that amygdala connectivity was modified following Pavlovian conditioning during the resting state^26^. However, many of these studies assessed brain activity during stimulus presentation using fMRI, without investigating post-reactivation processes^27,28^. A few studies evaluated brain activity during reactivation and extinction using EEG in a threat-conditioning paradigm, but only during stimulus presentation^29^. In this way, the present study provides some of the first evidence of post-reactivation processes in a threat-conditioning memory using EEG, through the analysis of brain activity in the time-frequency domain and by studying microstate dynamics.

In recent years, the use of Microstate analysis has significantly increased in the search for neural markers associated with various mental processes^30^. Microstates are short-lived, quasi-stable topographies of brain activity (lasting approximately 80–120 ms) that reoccur over time^31,32^. They have been proposed as fundamental building blocks of information processing in the brain^32,33^. Four canonical Microstates were defined based on their topography: Microstate A, B, C, and D^32^. These same Microstates have been consistently identified in a wide range of studies involving different cognitive processes^30^, and were also found in the present study. In particular, microstate C activity has been proposed to be related to mind wandering, with decreased activity being linked to externally focused attention^30^.

Building on these premises, the present study aimed to identify neural markers associated with the post-reactivation phase of threat-conditioned memories. Employing a three-day threat-conditioning protocol alongside EEG recordings taken during resting states before and after the reminder presentation, we investigated both oscillatory dynamics and Microstate organization to capture the rapid neural processes associated with memory reactivation.

This integrative approach provides new insights into the mechanisms through which reactivated threat memories are stabilized or modified, offering a framework for a deeper understanding of the neural basis of memory reactivation in relation to memory updating. This preliminary evidence suggests that adaptive updating mechanisms may provide new pathways to comprehend why, under certain circumstances, they fail to promote flexible regulation of aversive memories.

## Material and methods

Before the experiment, all participants provided written informed consent approved by the Ethics Committee of the Sociedad Argentina de Investigación Clínica (SAIC, Protocol #02/16), in accordance with the Declaration of Helsinki. A total of 52 participants between 18 and 35 years old were included (23 males and 29 females, mean age = 22.92 ± 0.54 years). Three other subjects were excluded from the analysis due to technical problems with the EEG recordings. Participants were randomly assigned to one of two groups: the Control group (15 females and 11 males, 23.03 ± 0.84 years) and the Reactivation group (14 females and 12 males, 22.08 ± 0.70 years).

### Subjective Assessment

The State-Trait Anxiety Inventory (STAI-S and STAI-T)^34^ and the Beck Anxiety Inventory (BAI)^35^ questionnaires were administered to assess participants’ anxiety and depressive levels, as these may affect fear conditioning, particularly threat conditioning^36,37^. Participants scoring above STAI > 45 or BAI > 30 were excluded; however, no participants were excluded based on these criteria.

### Experimental protocol

#### Reactivation Group

Participants assigned to the reactivation group completed a three-day consecutive experiment. On Day 1, they performed Stimuli Representation 1 followed by the threat conditioning. Twenty-four hours later (Day 2), a 4-minute resting-state EEG was recorded before stimulus presentation (Rest 1). After the incomplete reminder was presented (CS+ without the tone), a second resting-state EEG was recorded (Rest 2). On Day 3, memory evaluation (Test) was performed, followed by Stimuli Representation 2 (Figure 1).

**Figure 1.**
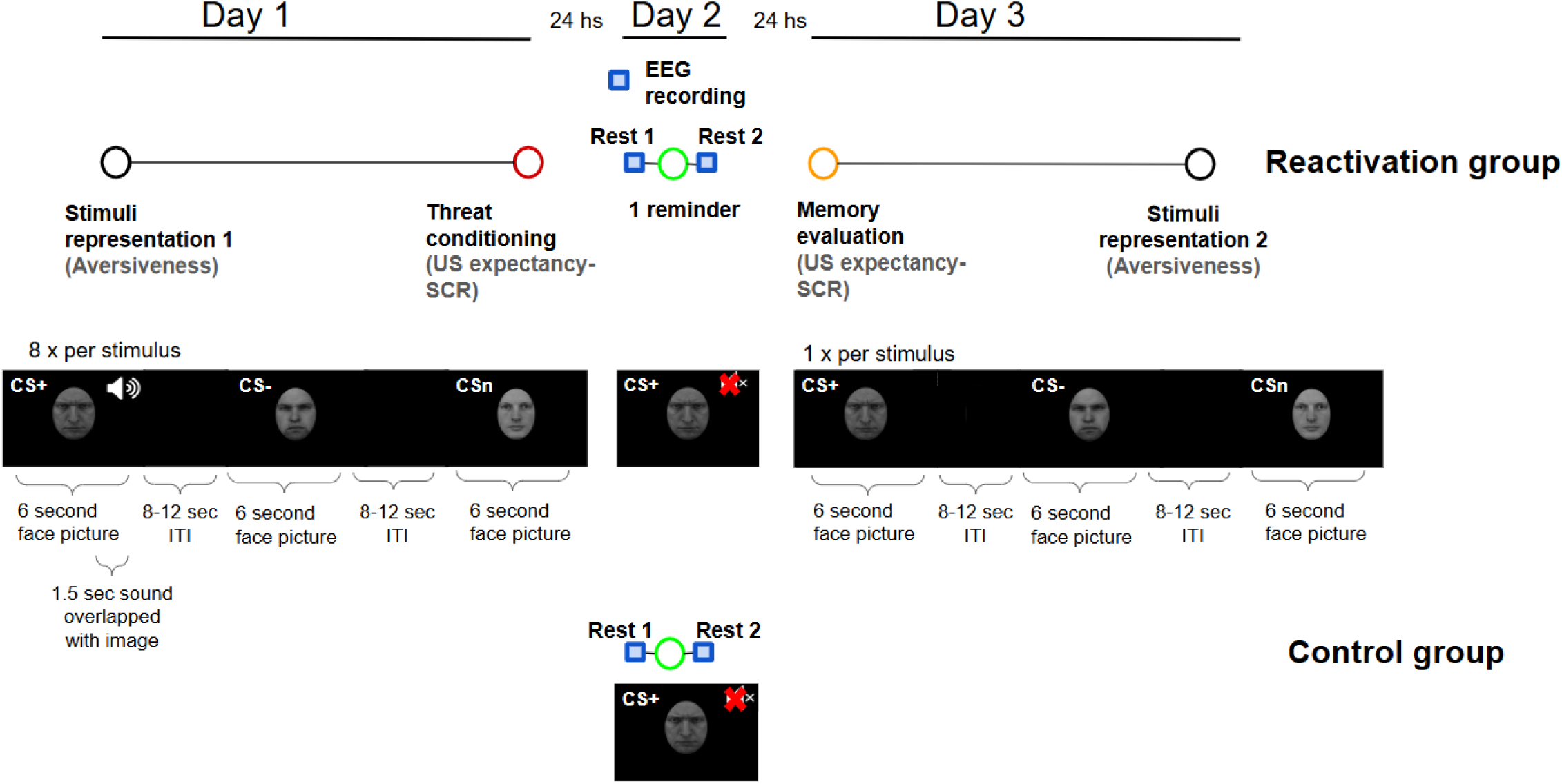
Schematic depiction of the experimental design illustrating group specifications across the three experimental sessions. Chronological outline of the experimental schedule for the two groups: Reactivation Group: performed the tasks across all three days, with a reminder of the CS previously paired with the US presented on the second day. Control Group: performed only the second-day task, observing the same reminder, but for them, it was the first time encountering it.

#### Control Group

Participants in the control group only completed the Day 2 task: a 4-minute resting-state EEG was recorded before stimulus presentation (Rest 1). After the incomplete reminder was presented, a second resting-state EEG (Rest 2) was recorded (Figure 1).

### Cognitive Bias

Stimuli Representation: Participants rated the aversiveness of 12 different male faces (seven angry and five neutral) on a 0–8 scale. Pictures were presented randomly and centrally on a black screen (each measuring 9.5 cm × 7 cm) until a response was given on the keyboard. Stimuli Representation 1 was performed before conditioning on Day 1, and Stimuli Representation 2 was performed after conditioning on Day 3. Supplementary figure 1

### Threat Conditioning

Following a paradigm similar to Picco et al., 2022^22^, two aversive (CS+ and CS–) and one neutral (CSn) male faces were presented centrally on a black screen (each slide measuring 9.5 cm × 7 cm) as conditioned stimuli. CS+ and CS– pictures were counterbalanced across participants. The unconditioned stimulus (US) was a 1.5-second auditory tone delivered through stereo headphones, generated by a TG/WN Tone-Noise Generator (Psychlab) and digitally controlled to maintain a mean intensity of 98 dB ± 2 dB. For each participant, US intensity was individually set to evoke an unpleasant but not painful sensation.

Acquisition phase: On Day 1, each face was presented 8 times in a different order (see Supplementary Figure 2). Each trial consisted of a 6-second face presentation followed by a randomly assigned 8–12-second inter-trial interval. The CS+ was paired with the US (tone) on 75% of presentations, overlapping the last 1.5 seconds of the CS. CS– and CSn were never paired with the tone.

Reactivation phase: On Day 2, the CS+ face (previously paired with the tone) was presented as before, but at the time the tone would have occurred, a screen indicating the end of the experiment was shown, and no tone was delivered.

### Memory evaluation (Test): On Day 3, one trial of each face (CS+, CS–, CSn) was presented as in the acquisition phase, but without the tone. (Figure 1)

During acquisition and memory evaluation, US expectancy and Skin Conductance Response (SCR) were measured as described below. All faces used in this experiment were taken from the Karolinska Directed Emotional Faces database which is free for research use (<u>database</u>).

### Measurements

#### US Expectancy

Participants responded “YES” or “NO” using a two-button keyboard, indicating whether they expected the tone to follow the presented face. Responses were considered valid only if made within the first 4.5 seconds post CS presentation (time interval preceding the US presentation).

#### Implicit Response

Sympathetic arousal was assessed via the rapid phasic components of electrodermal activity, specifically Skin Conductance Response (SCR), which reflects the emotional and cognitive states triggered by the CS^38^.

This involuntary physiological response reflects the level of excitability induced by different stimuli. When a stimulus (CS+) is paired with an aversive stimulus (US), the presentation of the CS+ generates an anticipation of the upcoming US, which increases arousal and may lead to an enhanced SCR. This effect has been previously observed in threat-conditioning studies^21,22,39^. SCR was recorded using a Psychlab Precision Contact Instruments device with a 50 Hz sine excitation voltage (±0.5 V). Two Ag/AgCl electrodes (20 mm × 16 mm) were placed on the intermediate phalanges of the index and middle fingers of the non-dominant hand. A five-point moving average filter was applied to smooth the raw SCR signal.

SCR was calculated as the difference between a baseline (mean conductance during the first 0.5 seconds after CS onset) and the mean conductance between 0.5 and 4.5 seconds post-CS. This time window was selected based on previous studies^7,38^. Each participant thus had one SCR value per trial (27 total: Acquisition: 8 CS+, 8 CS–, 8 CSn; Test –:1 CS+, 1 CS–, 1 CSn). SCR values were standardized into z-scores by subtracting the participant’s mean and dividing by the standard deviation. This normalization procedure was applied to the final baseline-corrected measures.

Outliers in skin conductance responses were removed using an interquartile range (IQR) based criterion. Data were grouped by stimulus type and Trial, and Q1, Q3, and IQR were computed for each group. Observations falling outside the range defined by Q1 − 2×IQR and Q3 + 2×IQR were excluded from further analyses. To perform the repeated-measures ANOVA, two participants were removed from the analysis after outlier removal.

#### EEG preprocessing

Four minutes of eyes-closed resting-state EEG were recorded for each participant before (Rest 1) and after (Rest 2) stimulus presentation. Given that the most robust results were observed during the last 90 seconds of Rest 1 and the first 90 seconds of Rest 2, this time interval was used for subsequent analyses (Supplementary figure 3). EEG was recorded using an Akonic Bio-PC 30 cap-mounted tin electrode system (10/20 international system, Electro-Cap International Inc.), referenced to the average of the two mastoids, sampled at 256 Hz. Data were bandpass-filtered between 1 and 35 Hz (8th-order Butterworth filter) and re-referenced to the average of all channels. Ocular artifacts were removed using independent component analysis (ICA), and erratic or deviating channels were visually inspected and interpolated as necessary. Data were discarded if more than three channels required interpolation, resulting in a total of 26 participants per group after preprocessing.

#### Time-frequency analysis

Time-frequency decomposition was performed using 7-cycle Morlet wavelets spanning 3-30 Hz. For each participant, the Rest 2 data were normalized by dividing by the time-averaged Rest 1 data.

### Microstate Analysis

Microstate analysis was performed using the Microstate EEGLAB toolbox^40^. After preprocessing (filtering, ICA, interpolation), Rest 1 and Rest 2 data from both groups were combined to generate Microstate prototypes.

The Global Field Power (GFP) was computed at each time point as the standard deviation of the potentials across electrodes, providing a reference-free measure of the overall strength of the scalp electric field. Specifically, GFP was calculated as

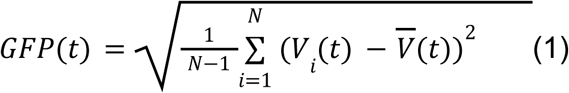

Where N signifies the total number of electrodes (channels), V_i_ (t) is the voltage recorded at electrode 𝑖 at time point 𝑡, and the average voltage across all electrodes at time point 𝑡.

Global Field Power (GFP) was calculated across all channels and frequencies between 1 and 35 Hz after preprocessing. Peaks of local GFP maxima (1000 peaks per participant, minimum 10 ms apart; peaks exceeding 1 standard deviation of GFP values were excluded) were used for clustering. A modified K-means algorithm with 50 random initializations optimized the number of clusters by minimizing the cross-validation criterion CV^40^.

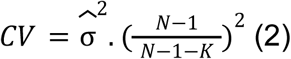

Where N signifies the Number of EEG channels, K the number of microstate clusters, an estimator of the variance of the residual noise.

Microstates were ordered by decreasing Global Explained Variance (GEV)^40^, where the correlation between each EEG sample (x) and its assigned prototype (a) was calculated

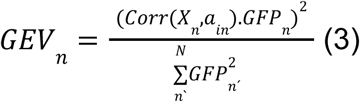

Microstates were labeled according to Tarailis et al., 2024: Microstate C (symmetric anterior-posterior configuration, GEV = 0.26), Microstate A (right frontal-left posterior, GEV = 0.15), Microstate E (posterior-central peak, GEV = 0.14), and Microstate B (left frontal-right posterior, GEV = 0.10). The optimal number of clusters was 4 (CV = 26.4, GEV = 0.64) (Figure 2).

**Figure 2.**
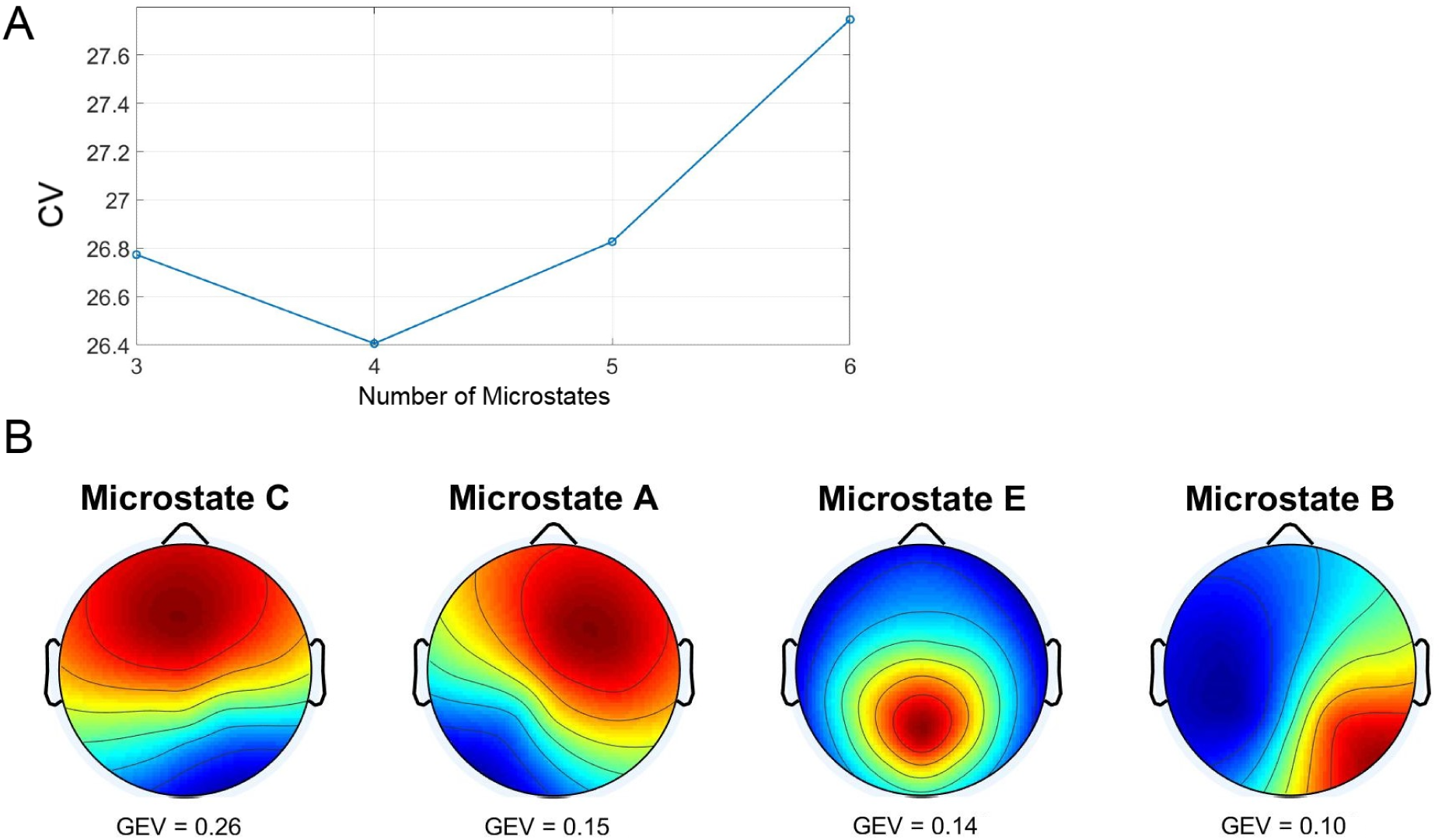
A) The CV was calculated for 3, 4, 5, and 6 Microstates, yielding an optimal number of 4 Microstates that minimized the CV. These are presented in B) Microstate prototypes ordered in decreasing GEV. They were labeled as C, A, E, and B according to Tarailis et al. (2024).

### Statistical Analysis

#### US Expectancy

Because including the participant as a random effect did not improve model fit, a generalized logistic regression model was used without random effects. Due to quasi-complete or complete separation in some conditions (all participants responding “yes” or “no”), odds ratios of responding “YES” were estimated using Firth’s penalized likelihood method (logistf package in R), which provides bias-reduced estimates and valid inference when standard logistic regression fails. Results are presented as odds ratios (ORs), comparing the probability of responding “yes” to each stimulus with the probability of responding “yes” to the CS+. OR values greater than 1 indicate that the likelihood of a “yes” response to the compared stimulus is higher than to the CS+. In contrast, OR values less than 1 indicate a lower likelihood compared to the CS+.

#### Implicit Response

To analyze the effect of stimulus type (CS+, CS–, CSn) on z-scored SCR for each trial, a repeated-measures ANOVA was conducted using R’s base aov function. The error structure accounted for the within-subject design (Error(Participant/stimulus)). Post hoc pairwise comparisons were performed using estimated marginal means (EMMs) with the emmeans package. Model assumptions were checked: residual normality (Shapiro-Wilk test and Q-Q plots), sphericity (Mauchly’s test via afex’s aov_ez function), and homogeneity of variances (Levene’s test from the car package). All assumptions were met (see Supplementary Table 1).

#### Cognitive Bias

Ordinal responses (Aversiveness scores) were analyzed using cumulative link mixed models (CLMM) via the ordinal package. Fixed effects included Time (Stimuli Representation 1 vs 2), Stimulus (CSs), and their interaction; participant was included as a random intercept. Estimated marginal means for interactions were computed with emmeans, and pairwise comparisons were adjusted with Tukey’s correction. Assumptions were tested: multicollinearity was assessed via Variance Inflation Factors (VIF < 5 acceptable), and the proportional odds assumption was tested using the nominal_test() function on a model without random effects. (see Supplementary table 2).

#### EEG Time-Frequency Analysis

Time-frequency data were compared between Rest 1 and Rest 2 within groups using non-parametric cluster-based permutation tests (FieldTrip toolbox)^41^. Monte Carlo methods with 1000 randomizations controlled for multiple comparisons by clustering electrodes with consistent differences. Two-sided dependent samples t-tests compared Rest 1 vs Rest 2 (cluster alpha = 0.05, minimum two neighboring channels per cluster, significance p < 0.05). Data were averaged over time. Between-group comparisons (Reactivation vs Control, Rest 2 normalized by Rest 1 baseline) were performed using independent t-tests.

Both analyses were conducted on the last 90 seconds of Rest 1 and the first 90 seconds of Rest 2.

#### Microstate Analysis

After deriving Microstate prototypes, statistics were calculated per participant, including Global Explained Variance (GEV), Global Field Power (GFP), duration, coverage, and occurrence. After removing outliers, these variables were analyzed per Microstate using linear mixed-effects models with Rest (1 vs 2), Group (Control vs Reactivation), and their interaction as fixed effects, and participant as a random effect. Significant interactions were followed up with post hoc tests using EMMs.

Model assumptions (normality and homoscedasticity) were checked via residual diagnostics (Q–Q plots, residuals vs fitted plots). No major deviations were found. Within the Reactivation group, participants were classified into two subgroups using k-means clustering (k = 2) based on standardized differences in evaluative scores and skin conductance responses (CS+ minus CS−) during testing, to compare Microstate measures between subgroups differing in their behavioral responses (Cluster 1 n=13; Cluster 2 n=12).

## Results

### US expectancy

The proportion of “yes” responses (indicating that the US was expected following the CS) was significantly higher for CS+ (Angry face paired with the annoying tone, US) than for CS− (Angry face not paired with the annoying tone) and CSn (Neutral face not paired with the annoying tone) starting from Trial 2 of the acquisition phase. This difference persisted throughout the acquisition phase and into the test phase.

During Trial 1, no significant difference was found in the proportion of “yes” responses to CS+ (46%) versus CS− (30%) (Odds Ratio (OR) = 0.53, 95% CI [0.17, 1.60], p = 0.26). However, a higher proportion of “yes” responses was observed for CS+ compared to CSn (0%) (OR = 0.028, 95% CI [0.0002, 0.25], p < 0.001).

By the end of the acquisition phase, in Trial 8, a marked difference emerged: 100% of participants responded “yes” to CS+ compared to 0% for both CS− and CSn (OR = 0.00035, 95% CI [0.0000011, 0.0073], p < 0.001).

A similar pattern was observed during the memory retention test phase: the proportion of “yes” responses to CS+ was 92%, compared to 7% for CS− (OR = 0.0065, 95% CI [0.0005, 0.041], p < 0.001) and 4% for CSn (OR = 0.0038, 95% CI [0.0002, 0.028], p < 0.001) (Figure 3a).

**Figure 3.**
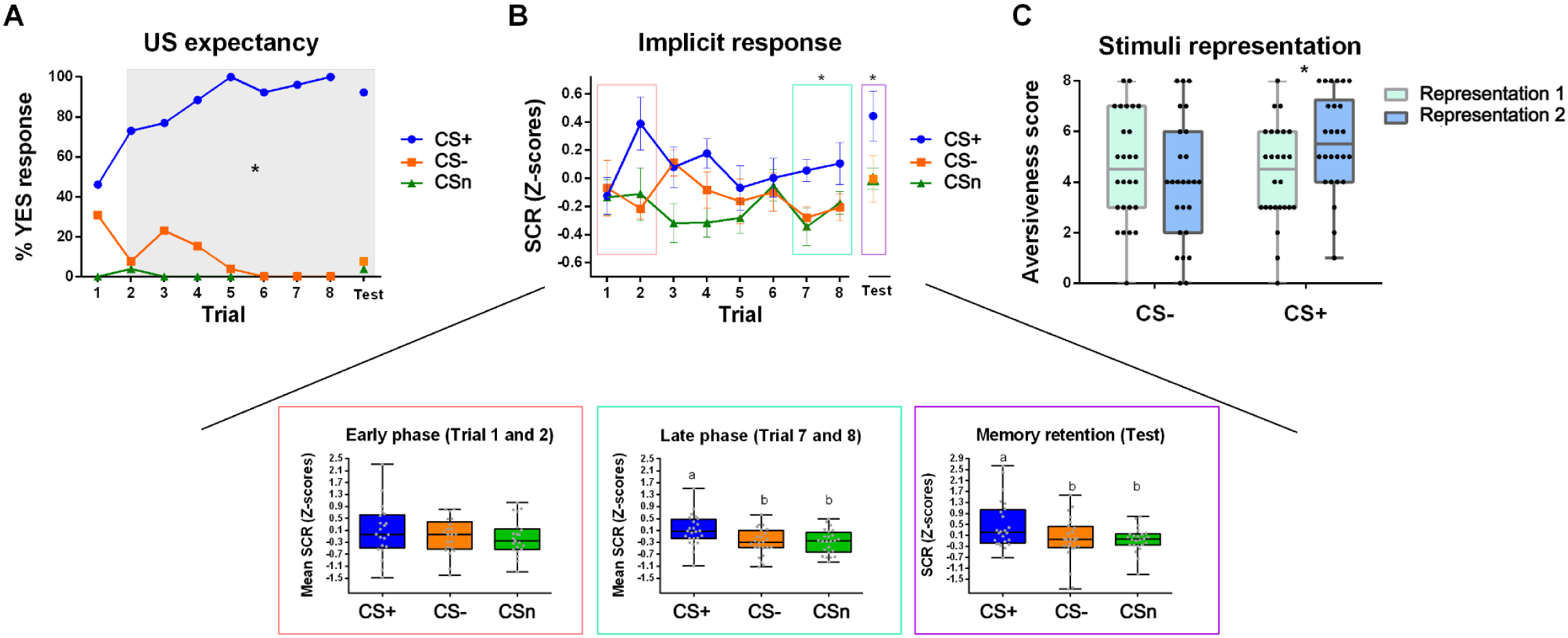
A) US expectancy represented as the proportion of “Yes” responses (“I will hear the tone after the CS”) for the three stimuli (CS+, CS–, CSn) across the eight conditioning trials and the test trial, 48 h later. B) Skin Conductance Response (z-scored) for the three stimuli (CS+, CS–, CSn) across the eight conditioning trials and the test trial, 48 h later. Boxplot of SCR (z-scored) for the early phase of conditioning (Trials 1–2), the late phase (Trials 7–8), and the test trial. Boxes represent the interquartile range, the line indicates the median, whiskers the minimum and maximum values, and each dot corresponds to each individual. Different letters indicate significant differences. C) Boxplot of stimulus ratings (aversiveness score) before conditioning (Rating 1) and after conditioning (Rating 2) for CS– and CS+. Boxes represent the interquartile range, the line indicates the median, whiskers the minimum and maximum values, and each dot corresponds to each individual. Asterisks indicate significant differences.

### Implicit response

A successful threat conditioning was observed, as evidenced by a greater skin conductance response (SCR) to the CS+ during the late phases of acquisition and the memory retention test, a difference that was not present during the early phases of acquisition (Figure 3b).

To evaluate the implicit response during threat conditioning acquisition, we compared SCR (Z-score) across stimuli (CS+, CS–, and CSn) during the early phase (Trials 1 and 2) and the late phase (Trials 7 and 8). We found that during the early phase, there were no significant differences between stimuli. However, by the end of acquisition, SCR was significantly higher in CS+ than in CS– and CSn, highlighting the effect of conditioning on sympathetic arousal.

Early phase (Trials 1 and 2): The main effect of stimulus was not significant, F(2, 46) = 2.54, p = 0.09. Figure 3b.

Late phase (Trials 7 and 8). A significant effect of stimulus was found, F(2, 46) = 5.66, p = 0.0063. Post hoc pairwise comparisons (adjusted for multiple testing): The estimated mean difference between CS1 and CS2 was 0.39 (SE = 0.13), t(46) = 2.96, p = 0.013; between CS1 and CS3 it was 0.38 (SE = 0.13), t(46) = 2.86, p = 0.016; and between CS2 and CS3 it was −0.01 (SE = 0.13), t(46) = −0.09, p = 0.995. Figure 3b.

To evaluate implicit memory retention, we compared SCR (Z-score) across stimuli (CS+, CS–, and CSn) on day 3 (Test). We successfully found memory retention, evidenced by the higher SCR (Z-score) for the CS+ compared with the CS- and CSn. A significant effect of stimulus was found, F(2, 46) = 4.38, p = 0.018. Post hoc pairwise comparisons (adjusted for multiple testing): The estimated mean difference between CS1 and CS2 was 0.53 (SE = 0.21), t(46) = 2.53, p = 0.038; between CS1 and CS3 it was 0.54 (SE = 0.21), t(46) = 2.59, p = 0.033; and between CS2 and CS3 it was 0.01 (SE = 0.21), t(46) = 0.052, p = 0.998. Figure 3b

### Cognitive bias

To evaluate cognitive bias associated with threat conditioning, we compared the aversiveness scores of the CS+ and CS- before and after conditioning. We found that after conditioning, only the CS+ was rated as more aversive/unpleasant; the aversive score of the CS- did not change, highlighting the effect of the association between that particular image (CS+) and the aversive tone on the aversive perception of the face.

The interaction between Representation (1 and 2) and stimulus (CS+ and CS-) was statistically significant (β = 2.41, SE = 0.75, z = 3.19, p = 0.001), suggesting that the effect of stimulus differed by representation. Neither the main effect of Stimulus (β = –0.34, p = 0.484) nor Representation alone (β = –0.86, p = 0.095) reached statistical significance. Meanwhile, no differences were found before and after conditioning for the CS- (Estimate 0.85, SE = 0.51, z = 1.67,n=26, p = 0.339). The aversiveness score was higher after conditioning for the CS+ (Estimate = −1.55, SE = 0.52, z = −2.96, n = 26, p = 0.015). Furthermore, the aversiveness score of the representation 2 was higher for the CS+ compared with the CS- (Estimate −2.06, SE = 0.55, z = −3.72, p = 0.001), a difference not found for the representation 1 (Estimate 0.34, SE = 0.49, z = 0.70, p = 0.89). Figure 3c.

### Neural correlates

Time-frequency analysis:

To evaluate changes in brain activity after the stimulus presentation, the power spectrum before (Rest 1) and after (Rest 2) presentation was assessed for all the channels, 3-30 range frequency, and time average (90 seconds).

In both experimental groups we found an increase in activity after the stimulus. Reactivation Group: n=26, p=0.0009 (cluster size of 247 data points). Control Group: n=26, p=0.0009. (cluster size of 371 data points). During Rest 2, the reactivation group showed increased activity across most channels, primarily in the 8–23 Hz frequency range; in the control group, it occurred between 13 and 30 Hz (Figure 4a).

**Figure 4.**
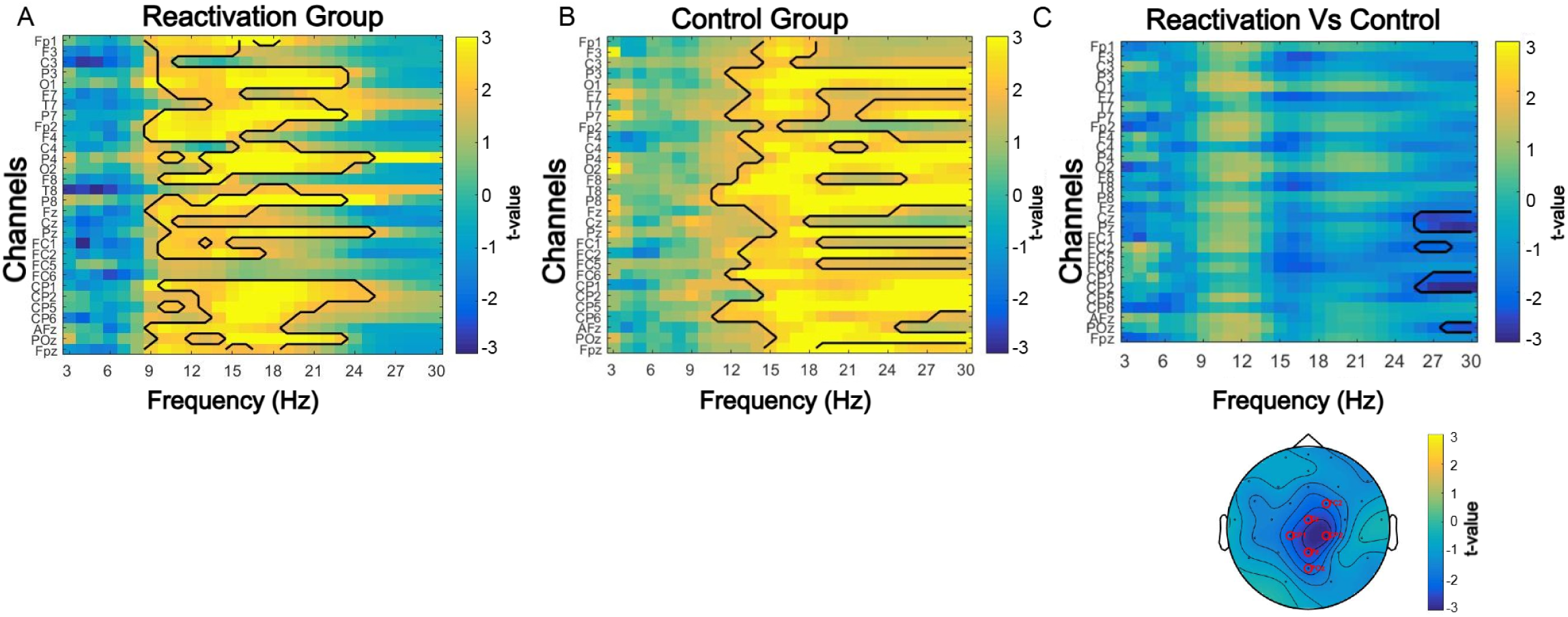
A) t-values from the cluster-based permutation test between Rest 2 and Rest 1 for the Reactivation and Control groups across all channels and frequencies (3–30 Hz). Contours indicate the significant positive cluster for each group. B) t-values from the cluster-based permutation test of Rest 2 (baseline-corrected to Rest 1) between Reactivation and Control groups, restricted to the frequencies and channels where a significant negative cluster was found in the between-group comparison. Topoplot shows the spatial distribution of significant channels and the corresponding t-values.

Based on this information, the next step was to compare post-stimulus brain activity (Rest 2, with Rest 1 as baseline) across conditions within the frequency ranges of interest (8–13 Hz and 25–30 Hz), as these bands showed different results across the experimental groups. An increased oscillatory activity in the control group relative to the reactivation group was identified over central regions in the 25-30 Hz frequency range (n=26, p=0.019, cluster size of 27 data points), Figure 4b. No differences were found for the 8-13 Hz frequency range.

### Microstates

After finding differences in the oscillatory brain activity between the experimental groups in the power spectrum analysis, we aimed to assess the effect on the level of transient changes in neural activity states by conducting a Microstate analysis.

#### Global Field Power (GFP)

The GFP represents the standard deviation of the voltage amplitude, providing an index of the strength of global activation associated with a given Microstate^30,40^. In the present study, we found higher GFP values during Rest 2 in the control group compared to the reactivation group across all four proposed Microstates.

Pairwise comparisons of estimated marginal means revealed that Control was associated with significantly higher GFP values than Reactivation for Microstates C, A, E and B (Microstate C: estimate = 1.12, SE =0.52, t(58) = 2.14, p = 0.036; Microstate A: estimate = 1.07, SE =0.48, t(57) = 2.23, p = 0.029; Microstate E: estimate = 1.17, SE =0.52, t(58) = 2.24, p = 0.028; Microstate B: estimate = 1.22, SE =0.43, t(61) = 2.82, p = 0.0065). Figure 5a

**Figure 5.**
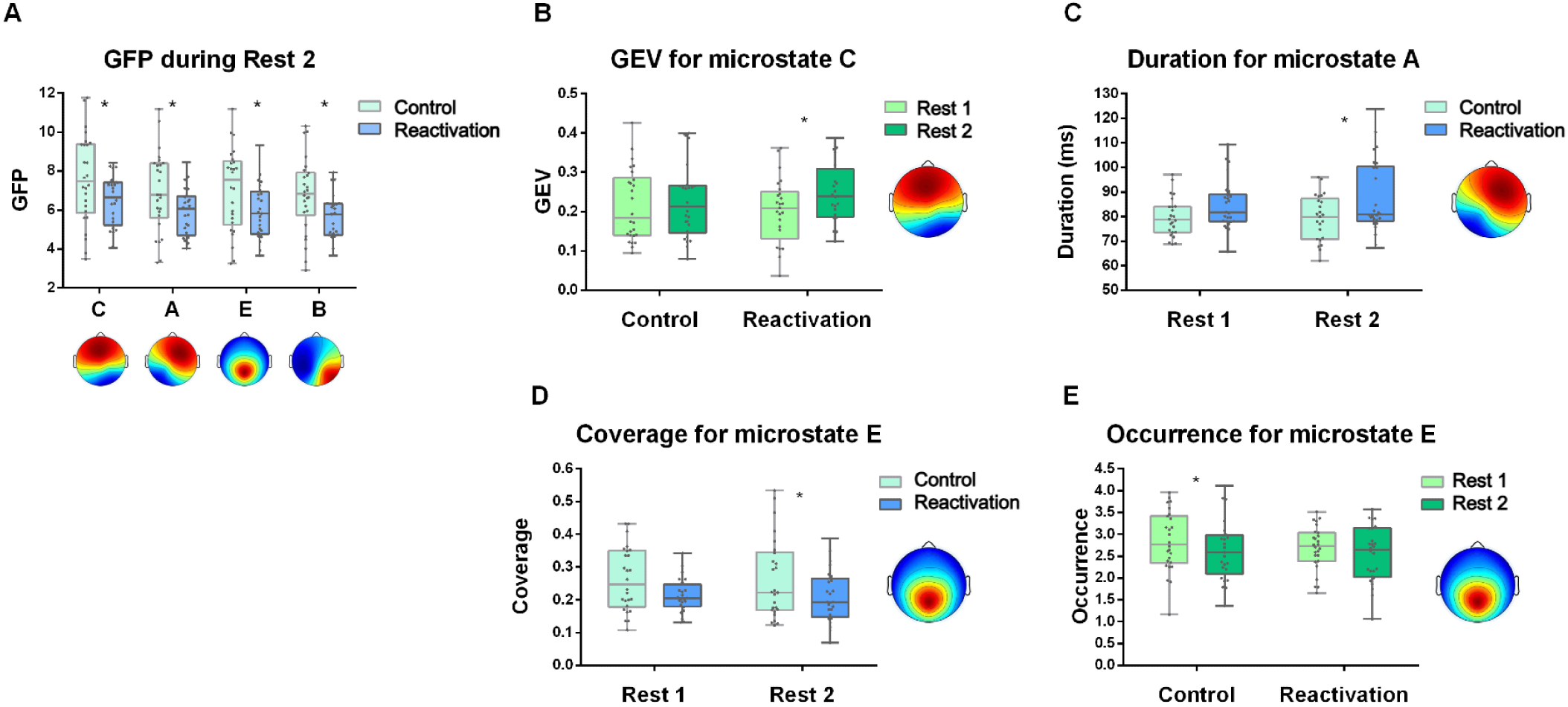
Boxplots showing interquartile range (boxes), median (line), minimum and maximum (whiskers), and individual values (dots) for: A) GFP for the 4 Microstates identified in the Reactivation and Control groups. Asterisks indicate significant differences between experimental groups. B) GEV for Microstate C comparing Rest 1 and Rest 2 in both experimental groups. Asterisks indicate significant differences between resting states. C) Duration (ms) for Microstate A in each experimental group during Rest 1 and Rest 2. Asterisks indicate significant differences between experimental groups. D) Coverage for Microstate E in each experimental group during Rest 1 and Rest 2. E) Occurrence for Microstate E comparing Rest 1 and Rest 2 in both experimental groups. Asterisks indicate significant differences between experimental groups and between resting states, respectively.

No differences were found during Rest 1 (see Supplementary Table 3)

#### Global Explained Variance (GEV)

The Global Explained Variance (GEV) indicates how well a given prototype explains the data. It is computed as the squared correlation between a sample and its assigned Microstate, weighted by the GFP at that specific time point^30,40^. We found an increase of GEV during Rest 2 only for Microstate C in the Reactivation group (Control group: estimate = −0.011, SE = 0.01, t(48) = −1.12, p = 0.268; Reactivation group: estimate = −0.03, SE =0.01, t(49) = −2.76, p = 0.008). Figure 5b.

#### Duration

Duration is defined as the average time a given Microstate remains active, expressed in milliseconds^30,40^. Significant differences were found only for Microstate A, with the Reactivation group showing longer durations during Rest 2, a difference that was not observed during Rest 1 (Rest1: estimate = −4.49, SE = 3.02, t(73) = −1.48, p = 0.141; Rest 2: estimate = −8.5, SE =3, t(72) = −2.83, p = 0.006). Figure 5c

#### Coverage

Coverage is defined as the fraction of time a given Microstate remains active^30,40^. Significant differences were found only for Microstate E, with the Reactivation group showing a lower coverage during Rest 2 compared with Control group, a difference that was not observed during Rest 1 (Rest1: estimate = 0.042, SE = 0.026, t(59) = 1.64, p = 0.105; Rest 2: estimate = 0.052, SE =0.026, t(58) = 2.03, p = 0.046). Figure 5d

#### Occurrence

Occurrence reflects the average number of times per second a Microstate is dominant^30,40^. We found a decreased occurrence during Rest 2 only for Microstate E in the Control group (Control group: estimate = 0.19, SE = 0.086, t(50) = 2.22, p = 0.03; Reactivation group: estimate = 0.155, SE =0.084, t(49) = 1.83, p = 0.072). Figure 5e.

Results from the other Microstates were present in supplementary table 4.

#### Relation between memory retention measures and brain activity

To evaluate whether brain activity differed according to responses to stimuli during the memory test, participants in the Reactivation group were divided into two subgroups based on behavioral variables: stimulus representation and SCR. Participants were clustered into two groups: one subgroup characterized as “good learners,” who showed an increased aversiveness score for the CS+ after conditioning and/or a greater SCR to the CS+ than to the CS– (Cluster 2), and another subgroup characterized as “poor learners,” who did not show an increase in CS+ aversiveness and/or did not exhibit a greater SCR to the CS+ (Cluster 1). Figure 6a.

**Figure 6.**
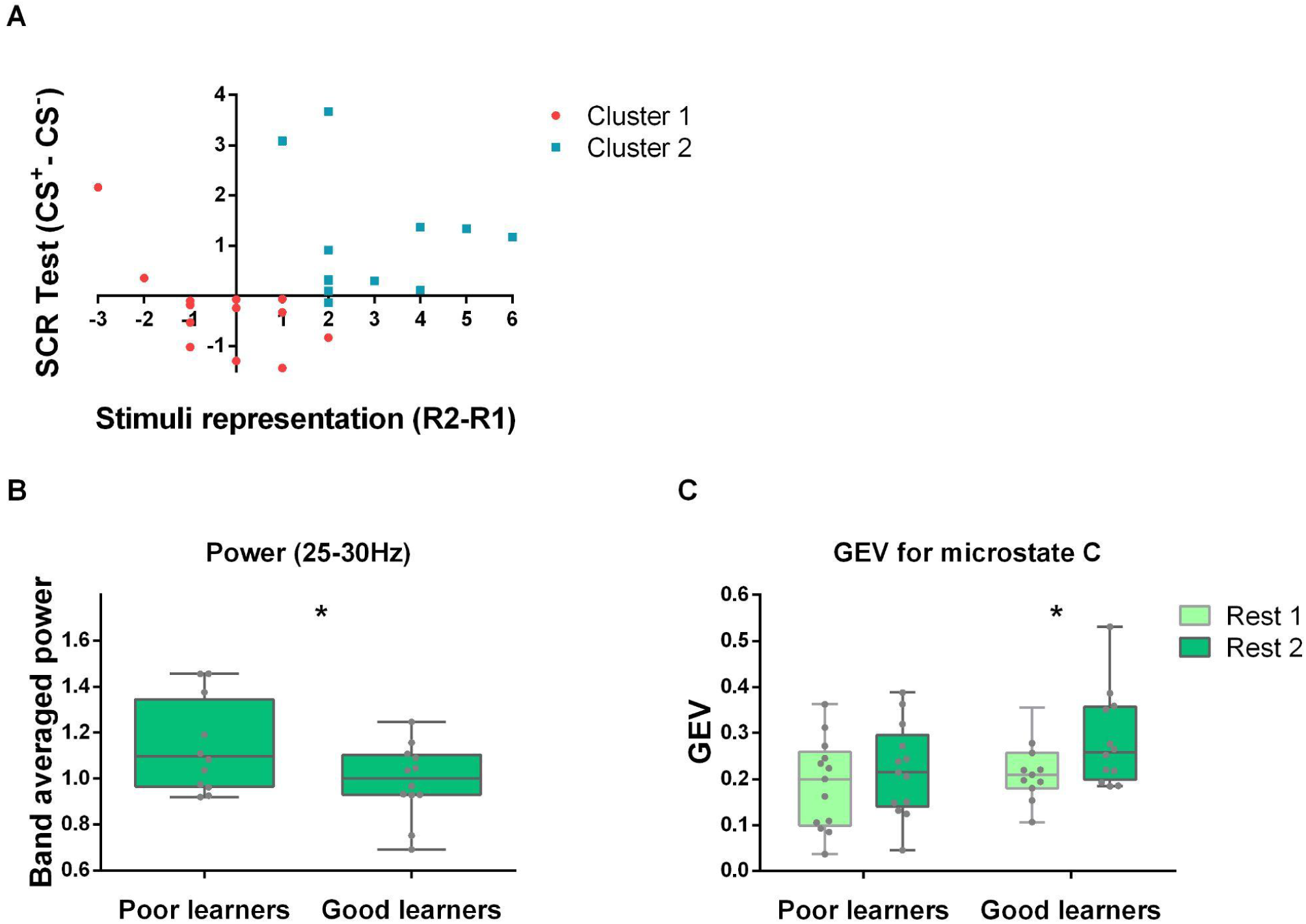
a) Participants were classified into two subgroups using k-means clustering (k = 2) based on standardized differences in evaluative scores and skin conductance responses (CS+ minus CS−) during testing. Cluster 1 was characterized as Poor learners, and cluster 2 was characterized as Good learners. b) Power for the significant electrodes found in figure 4C for 25-30 Hz, comparing Rest 2/Rest1 in Poor and Good learners. Asterisks indicate significant differences between resting states. c) GEV for Microstate C comparing Rest 1 and Rest 2 in Poor and Good learners. Asterisks indicate significant differences between resting states.

#### Power

As all assumptions were met (Shapiro–Wilk: p = 0.509; Levene’s test: p = 0.385), a linear model was performed to compare the power in Rest2, using Rest1 as baseline (25–30 Hz, averaged over time and restricted to the significant channel identified in Figure 4C). Poor learners showed greater power than good learners (t(22) = 2.10, p = 0.047). Figure 6b.

#### Microstate C

GEV increased after the reminder in good learners only, with no change observed in poor learners (poor learners: estimate = −0.028, SE = 0.022, t(23) = −1.3, p = 0.20; Good learners: estimate = −0.064, SE =0.024, t(24) = −2.58, p = 0.016). Figure 6c.

We found lower beta activity in the Reactivation group and an increased GEV for Microstate C (Figures 4 and 5). Similar results were observed when comparing the good learners subgroup with the poor learners subgroup, with the latter showing a pattern more similar to the Control group. We also found group differences in GFP that support these findings (Supplementary Table 5).

Results from duration, coverage and occurrence for all Microstates were present in supplementary table 6.

## Discussion

Adaptive behavior requires not only the detection of environmental threats but also the updating of previously acquired information. Memory reactivation facilitates this flexibility by making consolidated memories labile, allowing for modification through reconsolidation^14,17^.

The present study aimed to identify neural markers of the post-reactivation process of a threat-conditioning memory. We recorded EEG activity before and after reminder presentation in two groups: one that had undergone training (the Reactivation group) and another that had not (the Control group). We found a reduction in beta-band activity (25–30 Hz) in central brain regions and changes in the characteristics of microstate C after memory reactivation, which we therefore propose as neural markers of a post-retrieval process in a threat-conditioning memory.

As previously noted, the amplitude of brain oscillations can provide valuable insights into complex mental processes, with specific oscillatory frequencies associated with particular functions^23,42^. For instance, Zhu and colleagues utilized MEG to demonstrate a decrease in beta-band activity (15–25 Hz) during the reactivation of episodic memory^43^. Likewise, Griffiths and coworkers presented fMRI/EEG evidence suggesting that reductions in alpha/beta activity (8–30 Hz) are associated with enhanced information processing in associative memories^44^. These findings align with the desynchronization hypothesis, which underscores the importance of decreased 8–30 Hz activity during both memory encoding and retrieval^24^. According to this hypothesis, desynchronization is linked to a more nuanced representation of information, thereby facilitating both encoding and retrieval. Based on working memory studies in primates, Miller and colleagues have suggested that a decrease in alpha/beta activity may provide a framework for determining which brain regions are available or unavailable for activation, thereby regulating activity at other frequencies, such as the gamma band^45–47^.

In line with this framework, we found reduced beta activity (25–30 Hz) in the Reactivation group compared to the Control group, which could reflect greater post-reactivation processing of the memory, as opposed to participants without prior knowledge of the stimuli. Furthermore, considering that the image presented to the Reactivation group was previously paired with an aversive sound and could therefore trigger the recall of a threatening situation, the observed beta reduction is consistent with MEG results by Roxburgh and coworkers, who reported a decrease in the 14–30 Hz frequency band during periods of threat (unexpectedly delivered electric shocks)^48^.

Regarding alpha activity (8–13 Hz), we observed an increase in the Reactivation group that was absent in the Control group. However, no significant differences emerged between the two groups. Previous research has suggested that increases in alpha activity are linked to inhibition. Specifically, it has been proposed that

regions engaged in a task exhibit decreased alpha activity. In contrast, non-involved regions display increased alpha, thereby inhibiting potential interference and directing attentional resources^49^. In episodic memory interference paradigms, an increase in alpha/beta activity (11.5–20 Hz) has been observed in competing memory regions during retrieval, implying that such increases serve to inhibit interference from competing memories^50^. Given this evidence, the alpha increase seen in the Reactivation group may indicate an inhibition of interference with the reactivated memory. However, due to the lack of significant group differences, we cannot draw strong conclusions.

It is worth noting that many studies report decreases in alpha/beta and beta at the time of reactivation, mainly in episodic or working memory paradigms^24,51^. In contrast, our study assessed activity during 90-second resting-state intervals, which may reflect not only reactivation itself but also subsequent post-reactivation processes, specifically in the context of threat-conditioning memories. On the one hand, evaluating resting-state activity after reactivation is important for investigating post-reactivation processes, which remain relatively underexplored. Nevertheless, future studies should also assess brain activity during stimulus presentation in order to better distinguish neural activity associated with the reminder itself from subsequent post-reactivation processes.

EEG activity can be described as a dynamic system, with Microstates reflecting brief, stable topographies of electrical activity across the scalp. Resting-state EEG, widely used in clinical research, captures spontaneous neural dynamics that may reveal how large-scale brain networks operate^52–54^. Building on our spectral analysis of resting-state activity, we complemented this approach with Microstate analysis.

Four canonical Microstates (A–D) have been consistently described across a wide range of studies and cognitive domains, supporting their role as fundamental units of brain activity^32^. In line with this literature, we also identified these canonical Microstates in our data (Microstate D sometimes appears as an inverse topography with a posterior-central peak, as in the present work. Nevertheless, following the labels from Tarailis et al. (2024)^30^, we label these inverted Microstates as E). While Microstate analysis has been increasingly applied in psychiatric and neurological disorders, comparatively fewer studies have examined how these patterns relate to specific cognitive processes. In the present study, we aim to contribute evidence on Microstate dynamics following the reactivation of threat memories.

Microstate A has been primarily associated with auditory and visual processing; however, some evidence suggests it may also be linked to arousal^30^. Antonova and coworkers found a positive correlation between Microstate A duration and self-reported alertness during task performance, which also correlated with increased alpha activity (8–12 Hz)^55^. Consistent with this evidence, we found greater Microstate A duration in the Reactivation group, possibly reflecting a higher state of alertness due to the association between the CS+ and the aversive US.

Microstate E has been associated with emotional information processing and alertness^30,56^. In particular, its coverage has been negatively correlated with somatic awareness^57,58^. In our study, we found reduced Microstate E coverage in the Reactivation group, which may also reflect heightened alertness due to anticipation of the aversive sound. However, because the occurrence decreased only in the Control group, the effect of reactivation on Microstate E should be interpreted with caution.

Given that Microstate C accounts for the most significant proportion of total variance, we focused our analysis of behavioral correlations on this Microstate. Based on fMRI, EEG, and brain stimulation evidence, Microstate C has been proposed to reflect the default mode network (DMN)^30,59,60^ . The DMN is a network of interconnected brain regions, whose decreased activity has been linked to externally focused attention, whereas it is typically active during mind wandering or resting states^53,61,62^.

In our study, GFP was greater in the Control group across microstates. In particular, the reduced Microstate C activity observed in the Reactivation group may reflect increased attentional engagement with a salient stimulus. In contrast, the greater activity in the Control group may reflect mind wandering following the presentation of a stimulus without associated information. We also found that GEV increased after stimulus presentation only in the Reactivation group.

Interestingly, when we further subdivided the Reactivation group into two clusters based on their physiological responses (SCR) and cognitive biases, we identified “good learners” (those showing stronger CS+ than CS– responses after conditioning) and “poor learners” (those showing weaker or absent effects). In this case, GEV increased only in the good learners. We propose that, for good learners, the CS+ stimulus was likely more salient; in contrast, for poor learners, the CS+ was likely less salient, leading to brain activity patterns more similar to those of the Control group, for whom the CS+ carried no relevant information. These findings are further supported by the GFP analysis: poor learners showed increased post-stimulus GFP in Microstate C, possibly reflecting mind wandering, whereas good learners did not, consistent with a greater focus on the stimulus. Moreover, when comparing EEG power between these two subgroups, we found higher power in poor learners (averaged over the 25–30 Hz range and across the previously identified significant channels), similar to the lower beta activity observed in the Control group. This analysis allows us to evaluate differences in Microstates and power between groups separated by behavior, not only through implicit responses but also by incorporating cognitive biases.

We confirmed the successful conditioning and memory retention of our protocol through implicit and declarative responses, along with the associated cognitive biases. In particular, the CS+ stimulus (paired with an aversive tone, US) elicited greater skin conductance responses and higher aversiveness ratings compared to non-reinforced stimuli. These findings indicate that threat conditioning engages both physiological and cognitive responses, providing a solid framework for investigating how memory-updating mechanisms may facilitate or hinder the flexible regulation of aversive memories.

Previous results^22^ also demonstrated that presenting a reminder containing a prediction error 24 hours after acquisition induces memory labilization. Picco and colleagues found that when participants performed a high-demand cognitive task after a reminder presentation (CS+ without US), memory retention was weakened^22^. This effect was explained by competition for cognitive resources required for both the reconsolidation of the reactivated memory and the demanding task, ultimately weakening memory. By contrast, when participants performed a low-demand task after reactivation, no such memory impairment was observed. In line with these results, de Voogd and colleagues found that participants exposed to a working memory task during extinction showed reduced threat responses and enhanced extinction learning, which could be explained by competition between the salience and central executive networks of the brain^63^. These results confirm that the incomplete reminder used in this protocol elicits the reactivation of the conditioned memory and triggers post-reactivation processes involved in memory re-stabilization, which can be disrupted by an interfering task. Taking this into account, since EEG resting-state activity was measured after reminder presentation, the present study reflects brain activity associated with post-reactivation processes of

threat-conditioning memory. Nevertheless, the current study design does not allow us to disentangle the effects of the different components triggered by the reminder presentation. Future studies could evaluate brain activity during stimulus presentation in order to complement these findings with information related to neural activity specifically associated with stimulus processing itself.

Taken together, our results suggest that beta-band activity (25–30 Hz) in central regions and microstate C plays a crucial role in the post-reactivation processes of threat-conditioning memories. Furthermore, Microstate A appeared to be active for a longer duration following memory reactivation, and we propose a potential role for Microstate E. However, further studies are needed to clarify this. Finally, we suggest that the Microstate C pattern observed in the Control group reflects periods of mind wandering. In contrast, the association between this microstate and behavioral outcomes indicates that the reminder was more salient for participants who showed clearer evidence of learning and memory retention.

Limitation.

While our primary objective was to focus on the reactivation of threat memories, we cannot entirely rule out the possibility that episodic retrieval played a role, as the similarity of contexts across sessions might have prompted recall of the prior visit. This overlap could have influenced the neural patterns we observed. However, it is crucial to emphasize that the resting-state recordings were collected immediately following the presentation of the reminder cue. For the Reactivation group, this cue served as an implicit memory trigger, while for the Control group, it functioned as a neutral, meaningless stimulus. Despite the potential influence of episodic factors, the significant between-group differences in beta-band activity and microstate dynamics offer convergent evidence for distinct post-retrieval processes, reinforcing the notion that threat memories remain malleable even after reactivation. To address this issue, future studies could include an additional group in which participants perform a different task on day one in order to control for the previous laboratory experience without undergoing threat conditioning. Furthermore, evaluating brain activity during stimulus presentation, both during conditioning and reactivation, could strengthen our findings by allowing us to disentangle the specific effects of the reminder presentation from the post-reactivation processes.

Finally, evaluating the neural processes underlying the disruption of reconsolidation processes, for example, by assessing EEG resting-state activity before and after the Reactivation + HWM task, as well as the neural changes observed during the memory test associated with such disruption, could strengthen our findings by further clarifying the relationship between reconsolidation interference and the decrease in beta activity observed in the present study.

## Data Availability

Data from this study may be made available for all reasonable requests to the corresponding author.

## Additional Information

The authors declare no competing interests.

This work was supported by a grant from the University of Buenos Aires UBACYT 2023 (20020220400164BA) and CONICET PIP 2021-2023.

## Author contributions statement

L.B and M.E.P designed research; L.C performed research and analyzed data; L.B, M.E.P and L.C wrote the paper.

